# Cross-ancestry information transfer framework improves protein abundance prediction and protein-trait association identification

**DOI:** 10.1101/2025.08.13.670235

**Authors:** Wenli Zhai, Lingyun Sun, Wenwei Fang, Yidan Dong, Chunxiao Cheng, Yuanjiao Liu, Yuan Zhou, Jiadong Ji, Lang Wu, An Pan, Eric R. Gamazon, Xiong-Fei Pan, Dan Zhou

**Author notes:** Send correspondence to: Dan Zhou Xiong-Fei Pan; Eric R. Gamazon.

## Abstract

Genetics-informed proteome-wide association studies (PWAS) provide an effective way to map the complex molecular landscape of biological mechanisms for complex diseases. PWAS relies on an ancestry-matched reference panel to model protein expression using genetic variants as features and determine the protein’s impact on phenotype. However, reference panels from underrepresented populations remain relatively limited. In this study, we developed an analytic framework that borrows information from potentially multiple ancestries to boost the protein abundance prediction accuracy in an underrepresented population. We illustrate the framework’s utility and reproducibility through application to PWAS in East Asians: BioBank Japan (BBJ), Korean Genome and Epidemiology Study (KoGES), and Taiwan Biobank (TWB). An ensemble of information-sharing approaches was integrated to build the Multi-Ancestry-based Best-performing Model (MABM). MABM substantially improved the prediction performance with higher performance observed in both cross-validation and an external validation dataset (Tongji-Huaxi-Shuangliu Birth Cohort). Leveraging the BBJ, we identified three times as many significant PWAS associations with MABM as with the baseline Lasso model. Notably, 47.5% of the MABM specific associations were reproduced in independent East Asian datasets with concordant effect sizes. Furthermore, MABM enhanced gene/protein prioritization for downstream functional validation by (1) confirming a greater number of well-established gene/protein-trait associations and (2) identifying previously uncharacterized trait-associated genes. The benefits of MABM were further validated in additional ancestries and demonstrated in brain tissue-based PWAS, underscoring its broad applicability. Our findings close critical gaps in multi-omics research, develop a new reference resource of genetic models of protein abundance, and facilitate trait-relevant protein discovery in underrepresented populations.

## Introduction

The human proteome provides unique insights into human biology and disease^1,2^. As the final products of gene expression, proteins are subject to post-transcriptional and -translational regulation and processing, encoding additional functional information not detectable at the transcriptome level^3,4^. Case-control study using measured protein abundance is prone to reverse causality and confounding factors, while integrating information on genetic regulation may provide valuable mechanistic insights for discovery of potential druggable targets^5–7^. As both gene expression and protein abundance are fundamental molecular quantitative traits that may mediate genetic effects on phenotype, transcriptome-wide association studies (TWAS) frameworks^8,9^ have been extended to protein-wide association studies (PWAS). Relying on a reference panel, the PWAS methodology utilizes genetic variations as features to train protein abundance models, and apply the models to genome-wide association studies (GWAS) summary statistics to identify protein-trait associations^10–12^. However, optimal PWASs would require a well-powered reference panel with both genetic and protein data available as a training dataset, which remain limited in underrepresented populations. Models trained predominantly on European (EUR) reference panels often suffer from reduced predictive accuracy and increased false positive rates when applied to other populations^13,14^, limiting the applicability and statistical power of PWAS beyond EUR^15^. Therefore, accurate modeling across ancestries is essential to ensure generalizability of PWAS findings.

Early frameworks commonly relied on penalized linear regression to train imputation models for quantitative molecular traits. For instance, PrediXcan utilized an elastic-net model, which combines Lasso (L_1_) and Ridge (L_2_) regularization, to accommodate the balance between sparsity and polygenicity ^8^. Zhang et al. developed plasma protein elastic-net imputation models separately in European Americans and African Americans, and applied PWAS to serum urate and gout, revealing ancestry-specific genetic architecture ^12^. However, this study did not exploit the potential benefits of cross-ancestry information sharing, ignoring valuable potentially shared genetic regulation. To address this, several approaches have been developed to leverage multi-ancestry information. HEAT employs a reparameterized sparse group lasso to allow cross-ancestry information sharing while accommodating heterogeneous effect sizes and error variances^16^, but it is currently limited to two ancestral groups. Tian et al. proposed TL-Multi^17^, a multi-ancestry polygenic risk prediction method through transfer learning, which borrows useful information from auxiliary (European) data, improving learning accuracy for the target (non-European) data while maintaining efficiency and accuracy, although this method was not originally designed for imputation. Importantly, due to the complexity of the genetic regulation of protein abundance, a single regression model may only apply to certain patterns of regulation. A comprehensive methodology that integrates an array of empirically relevant models may lead to substantial performance gains in prediction performance and discovery power.

Here, we develop a PWAS framework, Multi-Ancestry-based Best-performing Model (MABM), that integrates an ensemble of statistical models for cross-ancestry protein abundance to enhance the prediction accuracy in an underrepresented population. To evaluate our approach, we compared the performance of MABM with the baseline Lasso model in an independent proteogenomic dataset, as well as the methods’ ability to identify protein-trait associations in several EAS GWAS datasets. Notably, the relatively larger sample size in a brain tissue dataset of EAS (compared to EUR) offers a unique opportunity to shift the conventional direction of information transfer in multi-ancestry studies and advance model performance in EUR. Finally, we showed the utility of MABM by mapping trait-associated proteins across multi-ancestry and evaluated its ability to validate well-established genes/proteins.

## Results

### Study overview

Here, we proposed a framework that improves the power of PWAS in an underrepresented population (e.g., EAS) by information transfer from other ancestries. Using the cross-ancestry UKB proteomics data, we trained genetics-based protein abundance prediction model in EAS using an ensemble of information-sharing approaches, which outperformed the baseline Lasso model in cross-validation. We leveraged the strengths of various models to select the best-performing model (referred to as MABM) for each protein and validated its performance in an external EAS dataset (Fig. 1a,b). We then applied MABM to the largest GWAS datasets to perform PWAS, including BBJ, KoGES, and TWB (Fig. 1c). We found that the enhanced prediction performance was reflected in an improved PWAS methodology using a number of benchmarking criteria: (1) number of identified protein-trait associations, (2) number and proportion of replicable associations across diverse EAS GWAS datasets, (3) number of well-established associations, (4) improved power in tissue-specific datasets (Fig. 1d), and (5) more accurate prioritization of likely causal genes at complex trait loci.

**Fig. 1:**
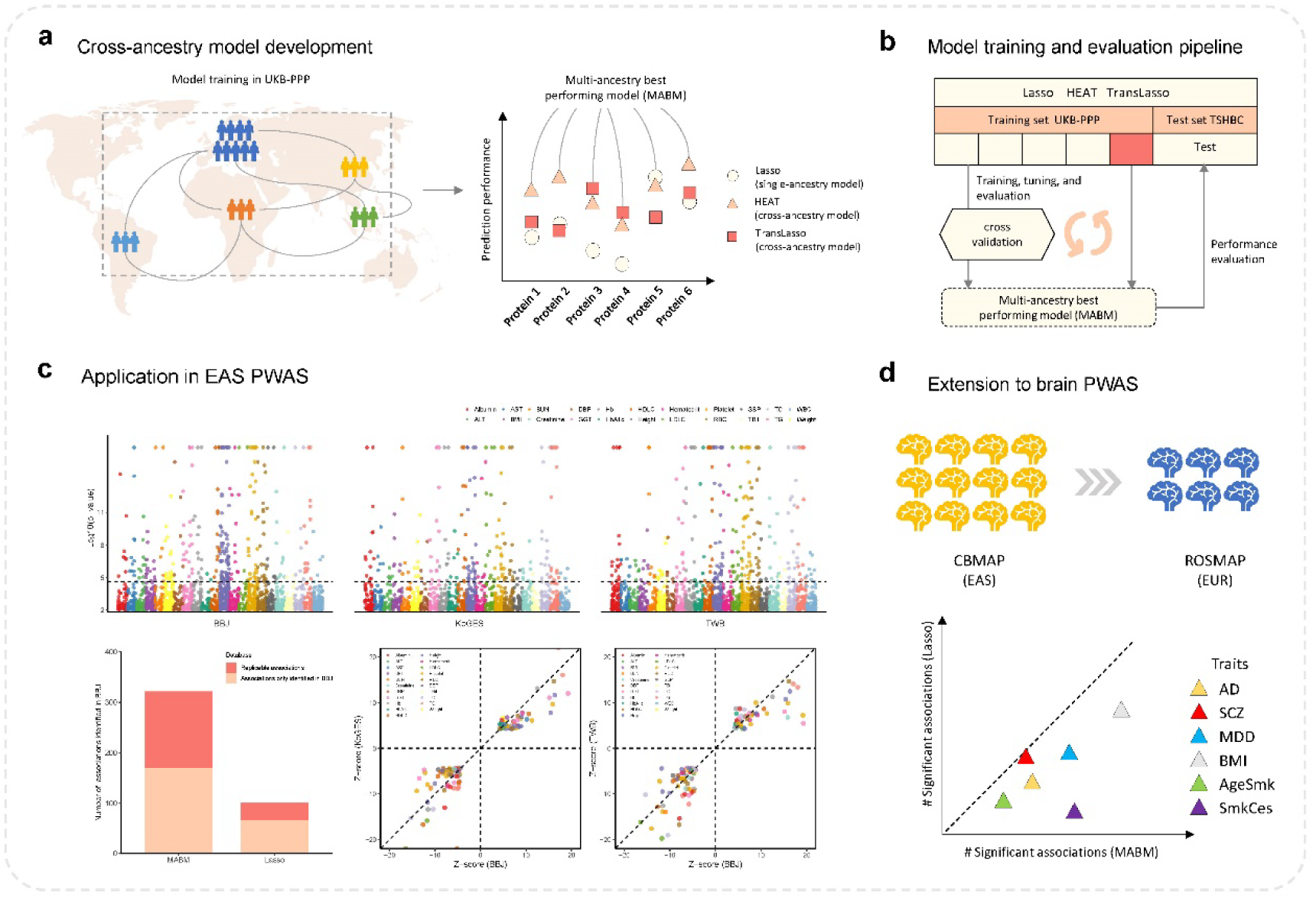
Overview of MABM model training framework and data analysis pipeline. **a** Borrowing information across ancestries using multiple model training approaches to build the multi-ancestry best-performing model (MABM) for modest-sized ancestries. **b** Illustration of model training, validation, and testing. **c** Application of MABM to East Asian GWAS data for protein-wide association studies (PWAS). **d** Bidirectional application of MABM enables improved PWAS in brain-specific European datasets with small sample sizes.

### Integrating multiple imputation models to improve EAS protein prediction

The EAS imputation models were trained using individual-level proteomic data from UKB-PPP. Specifically, Lasso was used to train prediction model solely on EAS ancestry data, while the HEAT model (based on sparse group Lasso) utilized both EAS and EUR data as training sets. TransLasso (derived from transfer learning) model was trained with EAS data as the target set and data from EUR, AFR, CSA, MID and AMR ancestries as the auxiliary sets. Both cross-ancestry models outperformed the baseline EAS-only Lasso model, increasing the number of iProteins (i.e., the number of models with predicted R^2^ > 1%) from 592 (Lasso) to 1,712 (HEAT) and 1,381 (TransLasso) (Fig. 2a,b). Particularly, for the less predictable iProteins (R^2^ < 5%) in the baseline lasso model, the median R^2^ improved from 2.27% (Lasso) to 3.88% (HEAT) and 4.14% (TransLasso). Given the possible range of genetic architectures of protein abundance, we noted that each approach had its strengths, theoretically and empirically. Therefore, we selected the best-performing model per protein (i.e., MABM), yielding 2,113 iProteins and further improving the median R^2^ from 2.12% (Lasso) to 3.93% (MABM) for low-predictability iProteins (R^2^ < 5% in the baseline Lasso model) (Fig. 2c). As illustrated in Fig. 2d, the cross-ancestry models (TransLasso and HEAT) predominately contributed to MABMs.

**Fig. 2:**
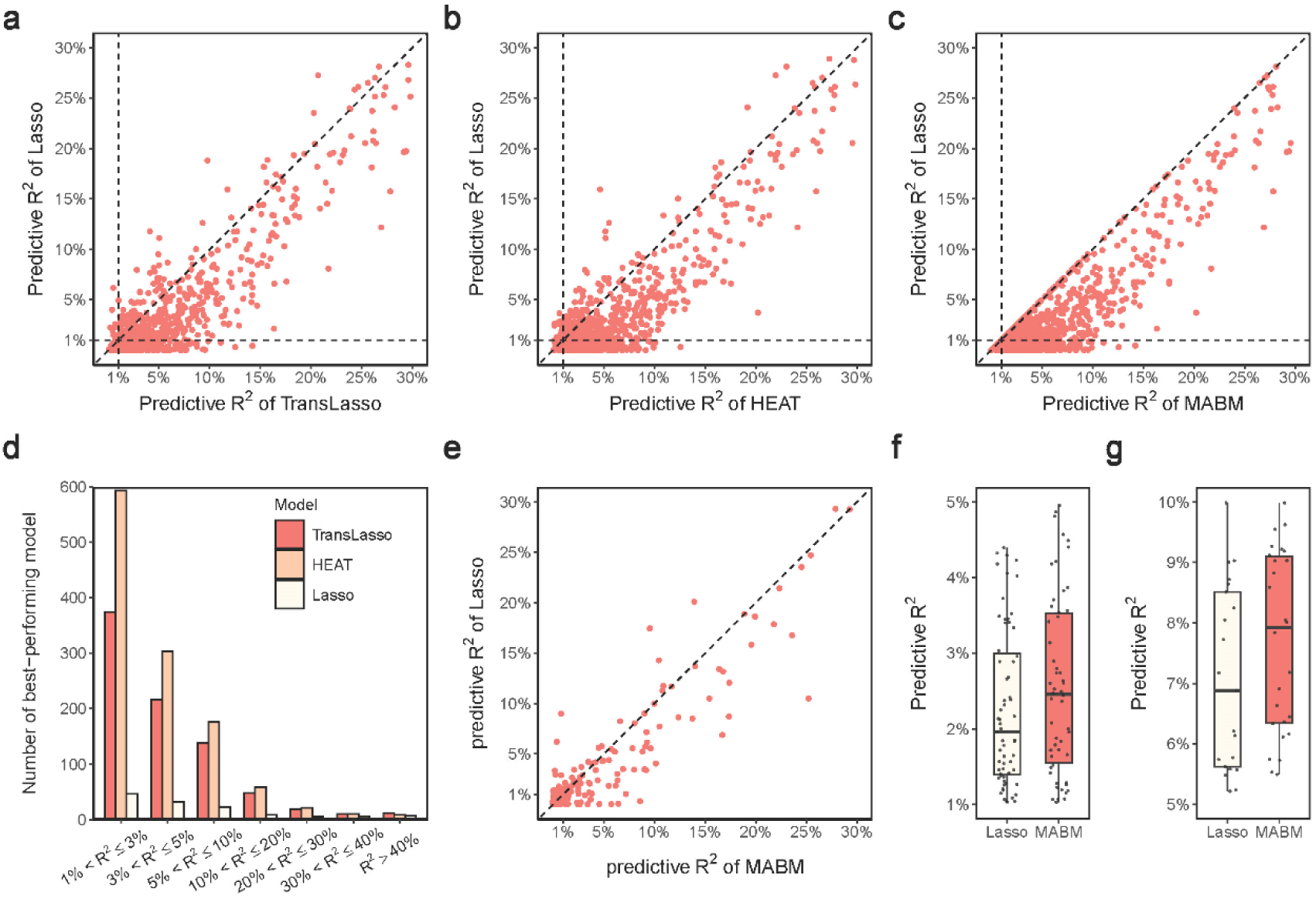
Performance comparison and validation of protein prediction models in EAS samples. **a-c** Cross-validation performance in the UKB-PPP EAS samples. Each point represents a protein. The predictive performance (R^2^) of TransLasso (**a**), HEAT (**b**), and MABM (**c**) is compared against the baseline Lasso model. The horizontal and vertical dashed lines indicate R^2^ = 1%; the diagonal dashed line represents y = x. **d** The composition of best-performing models (MABM) sourced from different model training approaches, partitioned by prediction performance R^2^. **e-g** External validation in THSBC. **e** Comparison of predictive R^2^ between MABM and Lasso for proteins with R^2^ > 1%; The predictive R^2^ for proteins that were not predictable by the Lasso model were scored as zero. **f** Boxplots of predictive R^2^ (1% < R^2^≤5%) for shared iProteins. **g** Boxplots of predictive R^2^ (5% < R^2^≤10%) for shared iProteins.

We validated the prediction performance in an independent EAS cohort (THSBC, *n* = 250). MABM successfully predicted 766 proteins, compared to 289 by Lasso, with 220 proteins shared across both models. Among the 220 shared iProteins, MABM significantly outperformed Lasso (paired t-test, *P*-value = 4.90×10^−6^, Fig. 2e). Among the 546 proteins uniquely predicted by MABM, 126 achieved a fair prediction quality (R^2^ > 1%). Notably, MABM showed consistent improvements for less predictable proteins (Fig. 2f,g): for iProteins with 1% < R^2^ ≤ 5%, median R^2^ increased from 1.96% (Lasso) to 2.46% (MABM); for those with 5% < R^2^ ≤ 10%, from 6.89% to 7.93%. Pairwise comparisons of the R^2^ of TransLasso, HEAT and Lasso can be seen in Supplementary Fig. 1.

### MABM identifies more protein-trait associations in the Biobank Japan

We applied MABM and the Lasso model to the GWAS summary statistics of 22 traits (Supplementary Table 1) derived from the BBJ GWAS dataset. At a Bonferroni-significant threshold of 5% (*P*-value < 2.38×10^−5^ for MABM and *P*-value < 8.45×10^−5^ for Lasso), 320 and 101 significant associations were found by MABM and Lasso model, respectively (Fig. 3). There were 65 protein-trait pairs that were significant in both sets of models. Notably, 255 unique significant associations were identified by MABM that were not found by Lasso, showing the substantial PWAS improvement in EAS ancestry. We repeated the same analysis in the KoGES, TWB, and an EAS meta-GWAS dataset (Supplementary Fig. 2 – Fig. 4) and summarized the results in Supplementary Table 2. Consistently, MABM identified more significant associations across all databases.

**Fig. 3:**
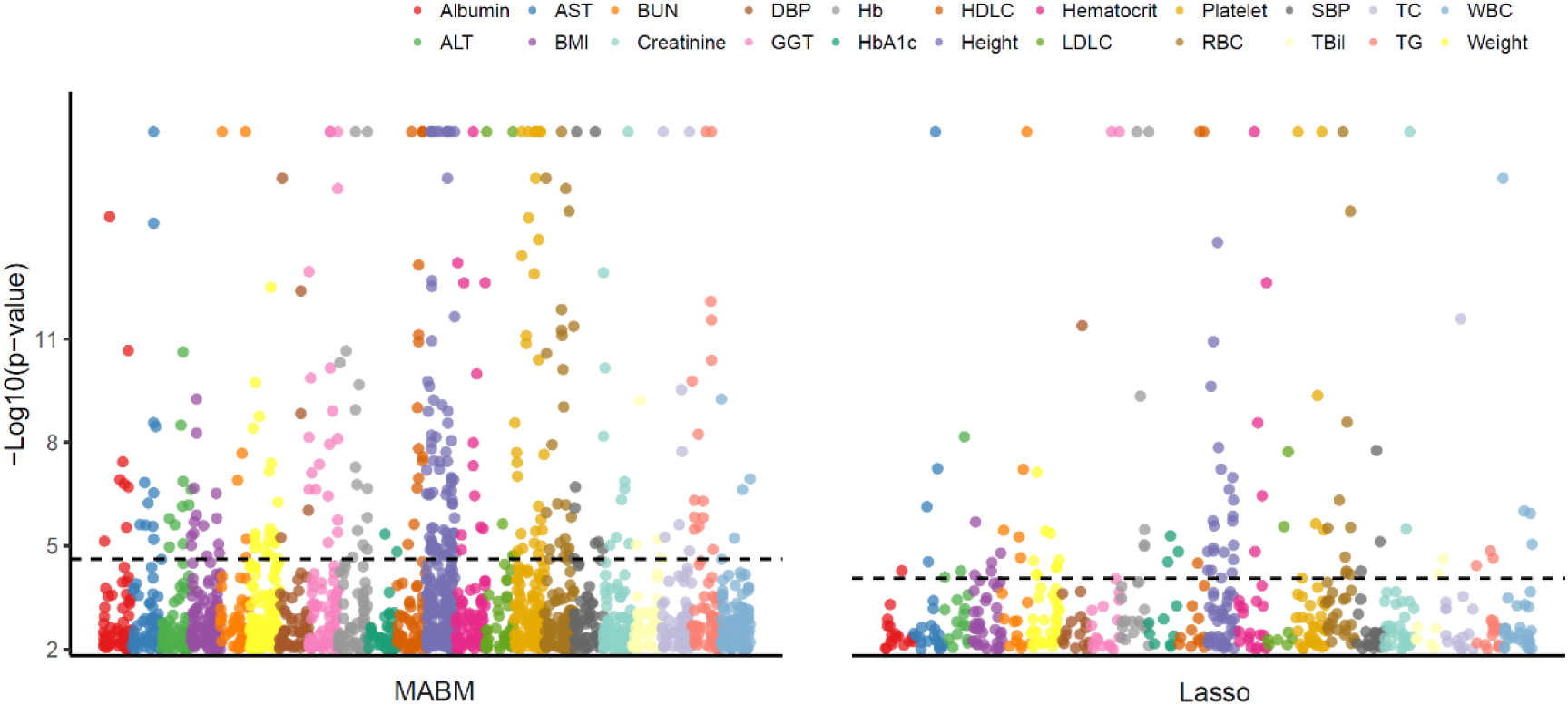
MABM substantially increased the identification of protein-trait associations. Based on the GWAS summary statistics from BBJ, PWAS was conducted for 22 complex traits using MABM (left) and the baseline Lasso model (right), respectively. The Manhattan plot shows the association signals. The dashed line represents the Bonferroni-significant level (*P*-value < 2.38×10^−5^ and *P*-value < 8.49×10^−5^, respectively) for the two models. The 22 traits of BBJ were labeled with distinct colors.

**Fig. 4:**
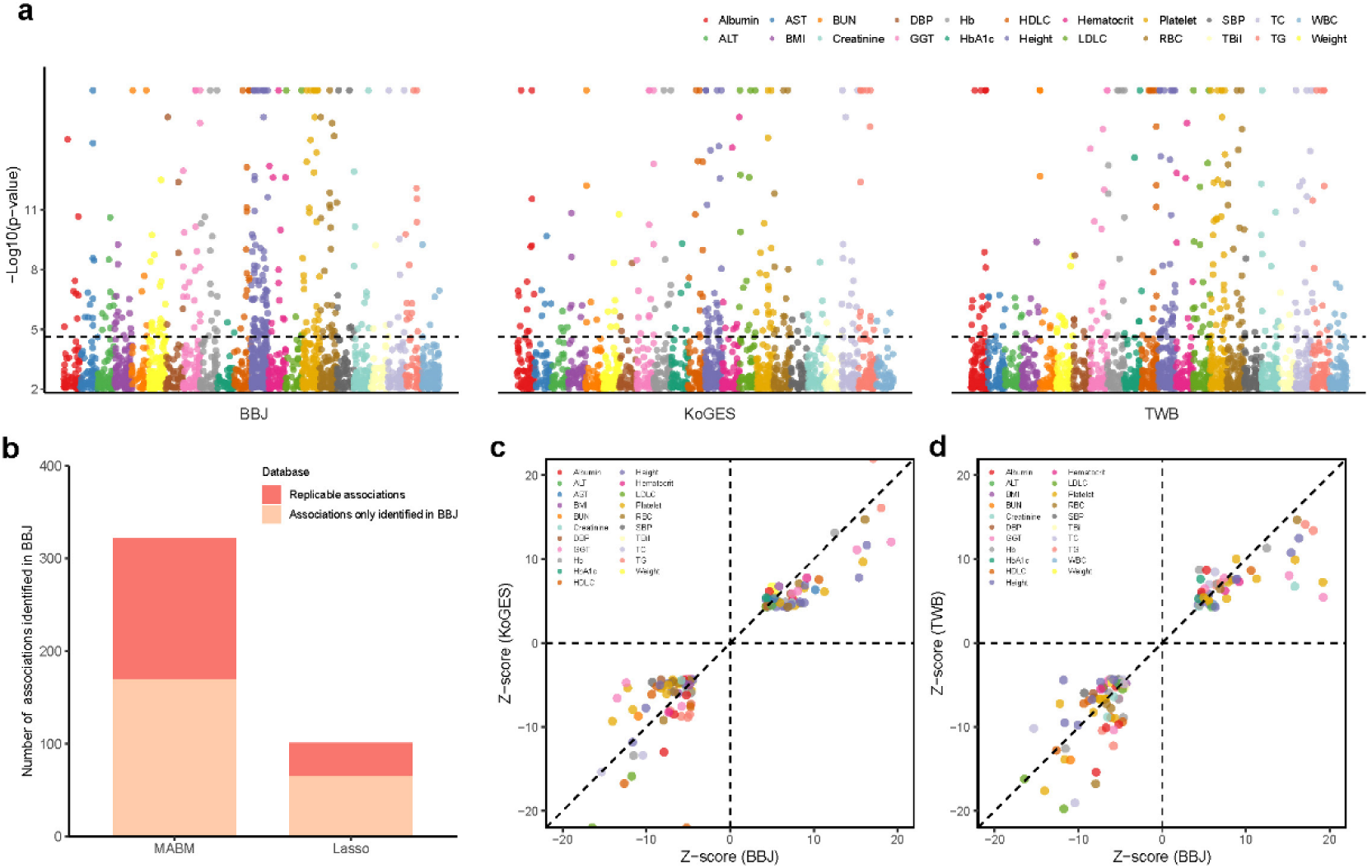
MABM identified replicable protein-trait associations across different GWAS datasets in EAS. **a** The Manhattan plots for the PWAS of 22 traits across three EAS datasets: BBJ, KoGES, and TWB. The horizontal dashed line represents the Bonferroni-significant level (*P*-value < 2.38×10^−5^). The 22 shared traits are labeled with distinct colors. **b** The number and proportion of significant associations from BBJ that are replicable in KoGES or TWB. **c-d** The consistency of association Z-scores (**c**) between BBJ and KoGES, (**d**) between BBJ and TWB. Each circle represents a significant protein-trait association pair that was identified by both GWAS datasets.

Additionally, we compared the performance of MABM and the Lasso model in identifying known gene/protein-trait pairs, using LDL-C and T2D as examples with 59 and 132 well-established genes/proteins, respectively (defined in Methods and Supplementary Table 3,4). Both sets of models were applied to LDL-C and T2D GWAS summary statistics in BBJ. For LDL-C, MABM identified five well-known genes/proteins (*ALDH2*, *SORT1*, *LDLRAP1*, *ANGPTL4*, and *LPA*) among 17 tested, while Lasso identified only one (*APOA2*). For T2D, MABM tested 16 genes and found significant associations for *GIPR*, *INSR*, and *GCG*. In contrast, Lasso tested only five genes and identified *GCG* alone. Comparison of PWAS P-values and five-fold cross-validated R^2^ (Table 1) showed that MABM yielded more accurate protein predictions than Lasso. Using the well-known genes as positive control (silver standard), overall, these results show that higher-quality imputations would facilitate the identification of trait-associated molecules.

**Table 1.** PWAS probes well-established gene/protein-LDL-C/T2D pairs by different prediction models.

| Trait | Gene | Model | $P$-value (MABM) | $P$-value (Lasso) | $R^2$ (MABM) | $R^2$ (Lasso) |
| --- | --- | --- | --- | --- | --- | --- |
| LDL-C | *ALDH2* | MABM | 0.0001 | -<sup>a</sup> | 0.0298 | -<sup>a</sup> |
| LDL-C | *SORT1* | MABM | 0.0000 | - | 0.0167 | - |
| LDL-C | *LDLRAP1* | MABM | 0.0073 | - | 0.0581 | - |
| LDL-C | *ANGPTL4* | MABM | 0.0374 | - | 0.0205 | - |
| LDL-C | *LPA* | MABM | 0.0004 | - | 0.0603 | - |
| LDL-C | *APOA2* | Lasso | 0.2451 | 0.0354 | 0.0361 | 0.0105 |
| T2D | *GIPR* | MABM | 0.0034 | - | 0.0151 | - |
| T2D | *INSR* | MABM | 0.0288 | - | 0.0127 | - |
| T2D | *GCG* | MABM /Lasso | 0.0055 | 0.0055 | 0.0198 | 0.0198 |

<sup>a</sup> Missing *P*-value/ $R^2$ denoted by dashes are due to either $R^2 \leq 1\%$ for the corresponding protein
abundance imputation models or no SNPs with non-zero weights in specific model due to variable selection.

### MABM identifies replicable signals across multiple datasets

We further analyzed the consistency of PWAS results obtained from different EAS GWAS datasets. Using MABM, PWAS was conducted on 22 common traits in BBJ, KoGES, and TWB (Fig. 4a), yielding 320, 207, and 269 significant associations (*P*-value < 2.38×10^−5^), respectively. Among them, 194 associations are significant in at least two datasets, with 83 associations being significant across all three datasets. We used the significant associations obtained in BBJ as a benchmark, counted the number of associations that can be reproduced in KoGES and TWB, and compared them with the signal reproducibility using Lasso. The results showed that 152 (47.2%) of the 322 significant associations based on MABM can be replicated in other Asian datasets, compared to 36 (35.6%) of the 101 based on Lasso (Fig. 4b). Among the 255 pairs derived from MABM from BBJ that were not found in Lasso, 97 (38.0%) and 90 (35.3%) associations can be reproduced in the KoGES and TWB, respectively, and 66 (25.9%) associations can be reproduced in all three datasets. The Z-scores of the reproducible significant protein-trait associations showed high consistency across BBJ and the other two datasets (Fig. 4c,d), with correlation coefficients of 0.952 and 0.947, respectively. Notably, all associations exhibit consistent effect directions.

Although we primarily demonstrated our method in EAS, it is also applicable to other non-European populations with limited sample sizes. We have included a parallel analysis for AFR and other ancestries in Supplementary results.

### MABM determines prioritization and fine-mapping of candidate genes by integrating European PWAS

Like TWAS, PWAS is susceptible to local co-regulation and LD contamination, often leading to multiple signals within a single locus. Given that many risk loci are shared across populations^18,19^, integrating PWAS results from both European and non-European ancestries may help improve fine-mapping resolution. Building on the PWAS signals observed in Europeans, we evaluated whether MABM-based PWAS in East Asians can better prioritize candidate genes compared to baseline Lasso models, using LDL-C and T2D as examples. Here, EUR protein imputation models were trained with Lasso using UKB-PPP data, and applied to GWAS summary statistics from UKB European samples. For LDL-C, we observed that MABM-based EAS PWAS successfully recovered five proteins (*SORT1*, *SDC1*, *LPA*, *NOTCH1*, *TRIM5*, *P*-value < 2.38×10^−5^) out of the 20 identified in EUR (*P*-value < 6.01×10^−5^, Fig. 5a), whereas the Lasso-based PWAS recovered only one (*NOTCH1*, *P*-value < 8.49×10^−5^, Fig. 5b). Notably, *SORT1* and *LPA* are established LDL-C risk genes^20^, while *SDC1*, *NOTCH1*, and *TRIM5* represent potentially novel associations supported across ancestries. Similar patterns were observed for T2D (Fig. 5c,d).

**Fig. 5:**
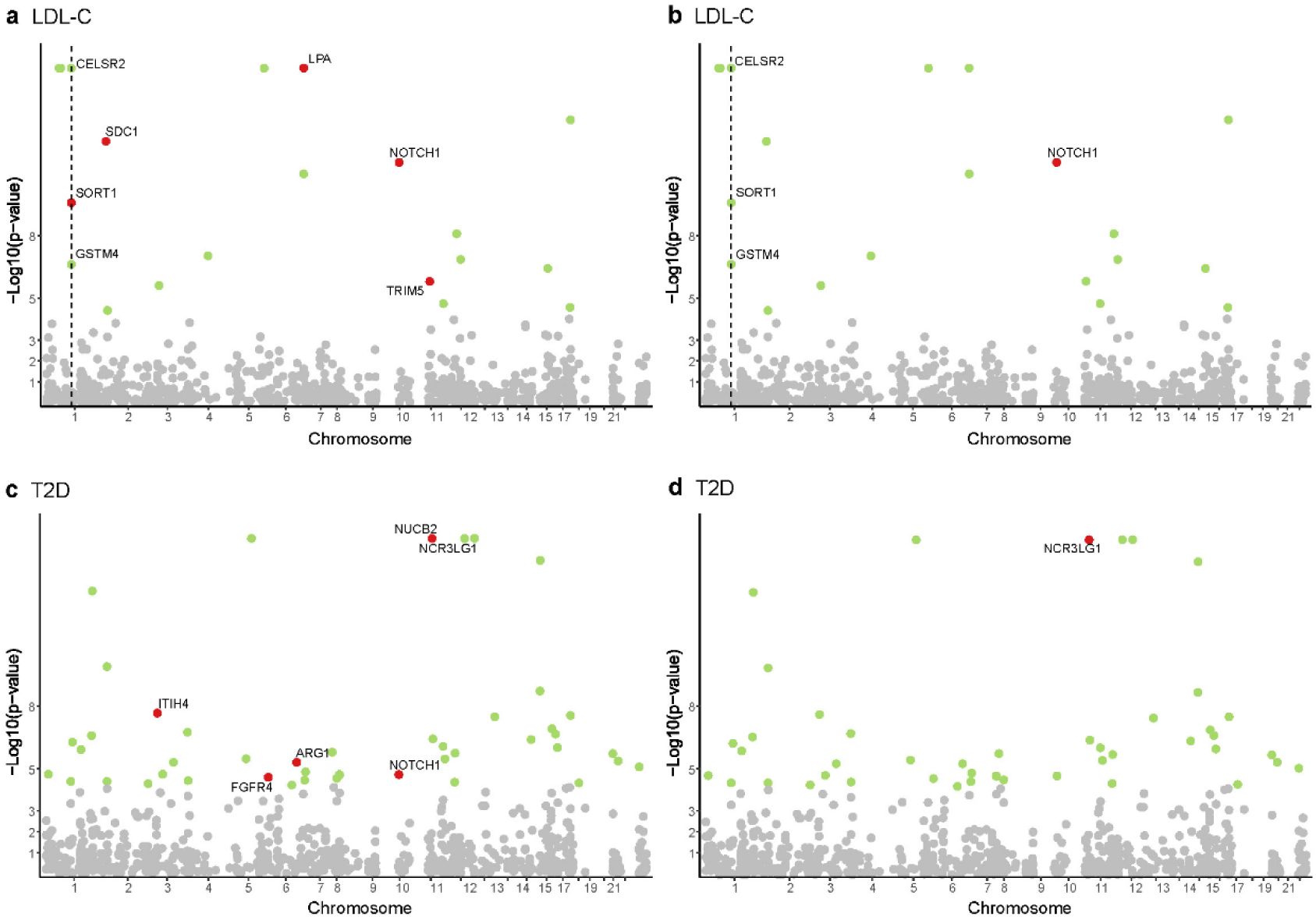
The integrated PWAS results across EUR and EAS. Panel **a** and **c** present the Manhattan plots of PWAS in EUR (**a**) for LDL-C and (**c**) for T2D. We further highlighted the results from MABM-based PWAS in EAS. Red, green, and gray circles denote proteins with both EAS and EUR statistical significance, only EUR significance, and EUR non-significance, respectively. The proteins with both EAS and EUR statistical significance are displayed using *P*-values in EUR PWAS and are labeled. Panel **b** and **d** present the same results except for the use of the baseline Lasso prediction model in EAS PWAS for LDL-C (**b**) and for T2D (**d**). The vertical dashed lines in **a** and **b** represent the position of *SORT1* on chromosome 1. *CELSR2* and *GSTM4* flanking *SORT1* are also labeled for illustrating the utility of EAS for prioritizing PWAS signals.

Moreover, we noticed that integrating East Asian and European PWAS results may further narrow the search space for potentially causal genes. For example, *SORT1* is a well-known LDL-C risk gene^21–23^, while multiple signals (*CELSR2* and *GSTM4*) have been observed in its flanking regions. In the PWAS of the European cohort, *SORT1* exhibited a modest signal compared to the stronger signal associated with *CELSR2* (Fig. 5a). The protein encoded by *CELSR2* is a member of the flamingo subfamily^24^, although the specific function of this particular member remains undetermined. Clearly, *SORT1* is more likely to be the pathogenic gene in this region. While EUR PWAS alone could not clearly prioritize *SORT1*, the EAS-based PWAS, particularly using MABM, provided additional support for its association with LDL-C. In contrast, the Lasso-based model failed to offer informative prioritization.

### Application of MABM to brain tissue data demonstrates flexibility across populations

To further evaluate the flexibility and generalizability of MABM, we applied it to proteomic data from brain tissue samples in two cohorts: ROSMAP (EUR, *n* = 318) and CBMAP (EAS, *n* = 729). In this analysis, which reverses the conventional direction of information sharing, we used the European ROSMAP samples as the *target* population, and leveraged the CBMAP data as the auxiliary ancestry group to build cross-ancestry MABM models. These models were then applied to nine brain-related traits using GWAS summary statistics from individuals of European ancestry (Supplementary Table 5).

In ROSMAP, we observed 3,747 and 2,925 iProteins (including 2,924 shared iProteins) were successfully predicted using MABM and Lasso, respectively (Fig. 6a). Among the 2,924 shared iProteins, MABM outperformed Lasso (paired t-test, *P*-value < 2.2×10^−16^). As a result, the median of predictive R^2^ increased from 1.94% (Lasso) to 2.45% (MABM) for the less predictable iProteins (R^2^ < 5% in the baseline Lasso model). Despite the limited sample size of the European brain proteomic reference panel, MABM identified substantially more significant protein–trait associations compared to the Lasso-based PWAS. Specifically, Lasso-based PWAS identified between 6 and 119 significant associations (27.4 on average), while the MABM-based PWAS identified between 17 and 150 significant associations, for an average of 56.1 (Fig. 6b and Supplementary Table 6). Among them, the largest improvement was observed for smoking cessation (SmkCes), where MABM uncovered 115 significant associations, compared to 17 from Lasso (6.8-fold improvement). We identified 17 genes significantly associated with at least three of the nine brain-related traits analyzed in the PWAS (Fig. 6c). Among them, *DDX39B*, *ATP6V1G2*, and *CLIC1* showed consistent associations with AD and multiple smoking-related phenotypes, including SmkCes, smoking initiation (SmkInit), and age at smoking initiation (AgeSmk). These findings raise the possibility that smoking behavior and AD may share common etiological pathways mediated by dysregulation of these proteins in the brain. The analysis illustrates the bidirectional utility of MABM for information transfer, demonstrating that cross-ancestry information can also be leveraged to improve discovery in EUR datasets with limited sample sizes.

**Fig. 6:**
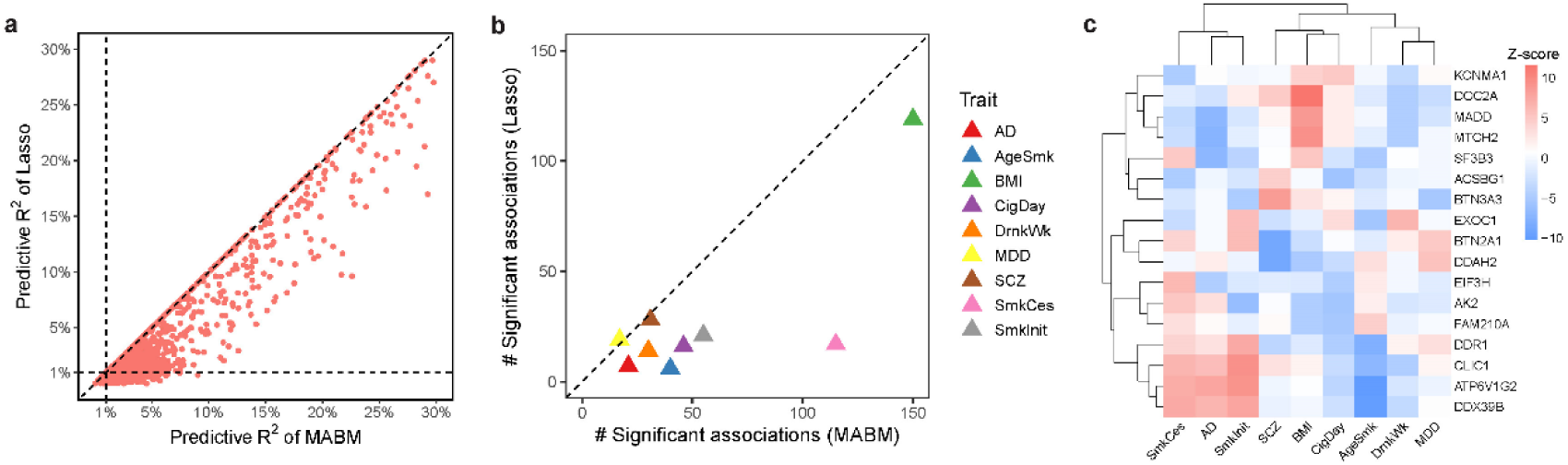
MABM exhibits strong performance in the ROSMAP brain proteomics data indicating strong bidirectional information transfer. **a** Comparison of predictive performance between MABM and Lasso using cross-validation in ROSMAP brain tissue samples. Each circle represents a protein. Horizontal and vertical dashed lines indicate the threshold of predictive R^2^ = 1%, with the diagonal dashed line (y = x) shown for reference. **b** Number of significant protein– trait associations identified by MABM (*P*-value < 1.37×10^−5^) versus Lasso (*P*-value < 1.72×10^−5^) across nine brain-related complex traits. Each triangle represents one trait and is labeled with a distinct color. **c** The heatmap displays MABM-based PWAS Z-scores for 17 genes (rows) across nine brain-related traits (columns). Genes and traits are hierarchically clustered based on Z-score profiles to highlight shared proteomic association patterns.

## Discussion

In this study, we developed an ensemble framework that captures the complexity of protein abundance regulation to enhance prediction accuracy and trait-associated protein discovery in ancestries or tissues with limited training sample sizes. By leveraging multiple modeling strategies—including TransLasso and HEAT, which borrow genetic regulatory information across ancestries—we showed substantially improved prediction performance compared with the baseline Lasso trained on single-population data. By adaptively selecting the best-performing models per protein, we demonstrated that MABM achieved higher predictive performance and identified more reproducible protein-trait associations in multiple underpowered contexts, including non-European populations and brain tissue. These results highlight the advantage of cross-ancestry information sharing in improving protein abundance prediction and association detection in cases where training data are limited.

As mentioned above, integrating cross-ancestry PWAS is a promising strategy for identifying potential pathogenic genes and proteins. European populations typically have larger sample sizes in GWAS, which enhance statistical power, increasing the ability to probe trait-associated genes/proteins^25,26^. However, studies that rely solely on European data may fail to adequately capture the genetic diversity of human populations, posing limited applicability for clinical applications and precision medicine^27–29^. Furthermore, linkage disequilibrium (LD) and pleiotropic effects may generate false positives, presenting a major challenge for result interpretation of both TWAS and PWAS^29,30^. Integrating multi-ancestry data may provide broader genomic coverage and more precise mapping for potentially causal biomolecules, offering insights for further prioritization of PWAS results and reducing the burden for functional studies. In practice, we successfully confirmed the well-known gene *SORT1* for LDL-C. Focusing on a specific locus at chromosome *1p13* which has the strongest association signals with LDL-C^31^, both *SORT1* and *CELSR2* showed associations in the European PWAS. Interestingly, PWAS in EAS based on MABM only supported *SORT1* which has clear evidence for lipid metabolism, while the PWAS based on the Lasso model failed to prioritize the proteins in this region. Calibrated by the ability to prioritize potentially causal proteins in the *1p13* region, we further show the effectiveness of boosting prediction quality using multi-ancestry data for model training.

We highlight some features of the framework. First, the framework can be extended to TWAS and other emerging omics association analyses in the future. Given that data from European populations are typically the first to be generated and often have sample sizes significantly larger than those from other populations, this framework is well-suited for such data. Second, similar to the Fusion framework^9^, MABM is capable of fitting a broader range of genetic architectures.

When the genetic regulation of the abundance of a protein is population-specific, lasso may perform better. Our framework can be reduced to this baseline model to optimize the selection of protein models. Moreover, the framework is designed to be flexible to incorporating additional prediction models in the future.

Our study has several limitations. First, most proteins have relatively low predictive R^2^ values in the model fitting of the first stage, possibly due to the low heritability of protein abundance driven by cis-SNPs. Trans-SNPs could be incorporated as additional features, potentially increasing performance, but this would come at the cost of increased noise and computational burden^32^. Second, the standard two-stage PWAS design does not distinguish between vertical and horizontal pleiotropy^33^. While cross-population analyses help to increase resolution for the search for truly pathogenic genes, true causality still requires validation through downstream biological experiments.

In conclusion, we developed a multi-ancestry-based ensemble framework, MABM, that aggregates multiple imputation models to enhance the accuracy of protein abundance prediction in a small reference panel. This framework can be easily applied to combine cross-ancestry reference proteogenomic datasets to identify SNP-trait associations in large-scale GWAS of complex polygenic traits. MABM substantially increases the number of replicable associations and provides additional insights into prioritizing potentially causal proteins.

## Methods

### The UKB dataset

The protein abundance prediction model was trained using the UK Biobank Pharma Proteomics Project (UKB-PPP, App No. 102158)^34^, which profiled 2,923 unique proteins from plasma samples of 54,219 participants using Olink Explore 3072 PEA. We retained 2,817 proteins with matched genotype data for 45,614 European (EUR), 931 African (AFR), 264 East Asian (EAS), 923 Central/South Asian (CSA), 311 Middle Eastern (MID), and 97 admixed American (AMR) individuals based on self-reported ancestry. Genotypes imputed with IMPUTE4^35^ were LD-pruned (R^2^≥0.8), and QC excluded individuals with >5% missing genotypes and variants with >5% missingness, MAF < 0.01, or HWE P < 1×10^−6^.

### The THSBC dataset

The protein abundance prediction models in the EAS population were evaluated in the Tongji-Huaxi-Shuangliu Birth Cohort (THSBC), which recruited Chinese pregnant women aged 18-40 years who attended their first prenatal check-up at Shuangliu Maternal and Child Health Hospital at 6-15 weeks of gestation. Women who had undergone assisted reproductive technologies or reported severe chronic or infectious diseases were excluded. This study was approved by the Ethics Committee of Tongji Medical College, Huazhong University of Science and Technology, Wuhan, China, and the reported investigation was conducted in accordance with the principles of the 2008 revised Declaration of Helsinki. Written informed consent was obtained from all participants before enrollment. After QC, 250 samples remained. Genotyping was performed using the Illumina Asian Screening Array (ASA), and 1,463 plasma proteins were measured using Olink targeted proteomics. Genetic variants were imputed using the Michigan Imputation Server (https://imputationserver.sph.umich.edu/index.html) with the EAS panel (1000 Genomes Phase 3 v5)^36^. SNPs with low call rates, HWE P < 1×10^−5^, or imputation quality ≤ 0.3 were excluded. A total of 1,445 overlapping proteins with UKB were retained for analysis.

### Brain tissue data

The China Brain Multi-omics Atlas Project (CBMAP) is a multi-omics study of human brain tissue based on a Han Chinese population, with participants sampled from northern, southern, and central regions of China^37^. Informed consent was obtained from donors or their legal representatives through voluntary donation agreements, allowing the use of both biological specimens and associated data for research. The study was approved by the Ethics Committee of Zhejiang University School of Medicine (Approval Nos. 2020-005 and 2024-007). Protein abundances in CBMAP were quantified using data-independent acquisition liquid chromatography–mass spectrometry (DIA LC-MS). A total of 10,911 proteins were identified, and 9,483 passed the <50% missing rate threshold. Intensity values were log-transformed, and missing values were imputed using k-means method. Genotyping quality control followed the same procedures as in the UKB-PPP, including LD-based pruning of SNPs with pairwise correlation ≥ 0.8. The Religious Orders Study and Memory and Aging Project (ROSMAP) is a longitudinal cohort focused on age-related neurodegenerative diseases, with participants recruited from religious communities and retirement facilities across the United States^38^ (access ID: 9603055). Proteomic data from ROSMAP were processed using the same pipeline as CBMAP. Finally, we analyzed 4,029 proteins and 216,416 SNPs shared by 729 samples from CBMAP and 318 samples from ROSMAP.

### Protein abundance imputation models

Protein abundance imputation models were trained using cis-SNPs (±500 kb around the transcription start and end sites of the corresponding protein-coding gene) as predictors and proteomic abundance as the outcome. The model weights of each protein were estimated using reference proteomics data. Assume that 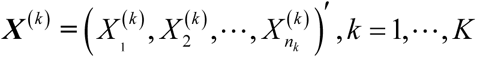 is the measured abundance level of a protein in *n_k_* samples of the *k*th ancestry, 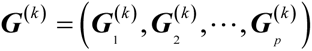 is the corresponding genotype matrix, and 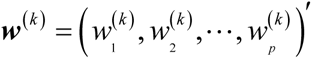 is the effect size to be estimated for the *k*th ancestry. We employed three methods—Lasso^39^, HEAT^16^, and TransLasso^40^—to construct the imputation models. The objective function of the Lasso training model can be expressed as:

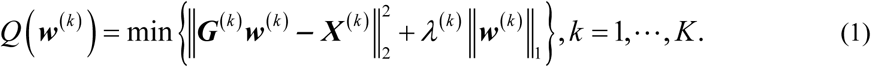

It is a linear regression model that uses L_1_ regularization, which drives some of the learned feature weights to zero, thereby achieving sparsity and enabling feature selection. Lasso was considered as the baseline model which was applied in PWAS using ancestry-matched EAS reference data for model training.

HEAT^16^ assumes that each ancestry has its own distinct regression coefficient vector and error variance:

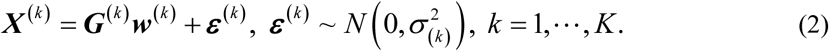

There are two main assumptions for HEAT. (1) Due to the sparsity of the genetic architecture of protein levels, where only a small number of SNPs contribute to the variance of protein levels, most entries in the weight matrix are assumed to be zero. Specifically, in the *k*th (*k* = 1, …, *K*) ancestries, the coefficient ***w***^(*k*)^ within the *k*th column of ***W*** = (***w***^(1)^, …, ***w***^(*k*)^)∈ℝ*^p^*^×^*^K^* is expected to be sparse, which is consistent with the sparse genetic architecture of protein levels; (2) Given the relatively high correlation of pQTL effect sizes across different ancestries, a considerable proportion of pQTL sites are assumed to be shared across ancestries. That is, for SNP*_i_* (i.e., the *i*th row of ***W***), the coefficients across the *K* ancestries tend to follow a pattern where they are either all-zero or with many non-zero entries across all ancestries. Consequently, HEAT incorporates a sparse group Lasso penalty:

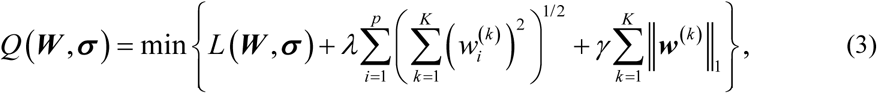

where 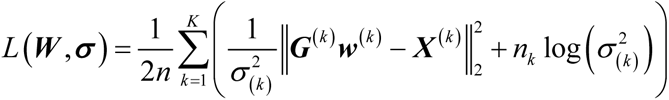 is the negative log-likelihood. The sparse group Lasso penalty models sparsity between and within groups in regression, enabling variable selection by incorporating prior grouping information of the features.

TransLasso^40^, a high-dimensional model derived from the idea of transfer learning, is designed to transfer knowledge between auxiliary and target samples, thereby enhancing learning performance of targets. Notably, TransLasso can incorporate information from multiple diverse auxiliary datasets, so it has application potential in cross-ancestry multi-omics studies. The target model of TransLasso for non-Europeans (i.e., EAS) can be written as

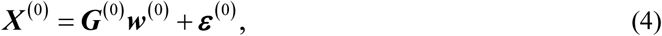

In the context of transfer learning, additional samples from *K* auxiliary ancestries are considered, i.e., *K* auxiliary models

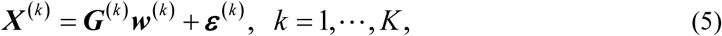

The similarity between the target model and given auxiliary models is characterized by the sparsity of their contrast vectors. In short, TransLasso first computes an initial estimator ***ŵ*** using the auxiliary samples. However, this estimator deviates from the target estimator ***ŵ***^(0)^ by a bias term ***δ̂*** = ***ŵ***^(0)^-***ŵ***. The target samples are then used to correct this bias. In fact, ***δ*** is a sparse high-dimensional vector with its L_1_-norm constrained to a relatively small range. The TransLasso algorithm developed by Li et al.^40^ is available to interested readers. Given the low statistical power due to small sample sizes from underrepresented ancestries, leveraging genetic knowledge from other ancestries is expected to significantly improve the estimation and prediction of protein imputation models for the target ancestry.

### Model training and performance evaluation

The optimal hyper-parameters of the training models were selected based on the cross-validation error, and the final models were refitted using all data with the selected parameters. Model performance was evaluated via nested 5-fold cross-validation (Supplementary methods). The imputable proteins with R^2^ > 1% were defined as iProteins.

### PWAS analysis by applying prediction models to GWAS datasets

We performed proteome-wide association studies (PWAS) by applying the protein prediction models (Lasso and MABM) to GWAS summary statistics using two-stage PrediXcan framework^8^. We primarily conducted PWAS for 22 phenotypes (Supplementary Table 1) shared across three East Asian biobanks—BioBank Japan (BBJ)^41^, Korean Genome and Epidemiology Study (KoGES)^42^, and Taiwan Biobank (TWB)^43^. We further used meta-GWAS summary statistics of TWB and BBJ to maximize the power of PWAS in the EAS population^44^. The GWAS information of the nine brain-related phenotypes is provided in Supplementary Table 5. To benchmark biological relevance, we focused on two well-characterized traits: low-density lipoprotein cholesterol (LDL-C) and type 2 diabetes (T2D). The LDL-C risk genes were collected from the KEGG cholesterol metabolism pathway and literature-based silver standards^20^, comprising 59 risk genes. T2D risk genes (*n* = 132) were curated by Mahajan et al. (https://t2d.hugeamp.org), categorized by strength of biological evidence. For both traits, BBJ GWAS summary statistics were used.

## Data availability

The Olink proteomics data from UKB-PPP are available under dataset https://biobank.ndph.ox.ac.uk/. The summary statistics for KoGES used in this study are downloaded from the KoGES Zenodo (https://zenodo.org/record/7042518), BBJ summary statistics from the Biobank Japan PheWeb (https://pheweb.jp/), TWB summary statistics as well as TWB and BBJ LDL-C meta-GWAS summary statistics from GWAS Catalog (https://www.ebi.ac.uk/gwas/publications/38116116), LDL-C and HDL-C summary statistics for Europeans from UK Biobank (http://www.nealelab.is/uk-biobank). The T2D ancestry-specific GWAS summary statistics are available through the DIAGRAM Consortium website (http://www.diagram-consortium.org/downloads.html). Summary statistics of GWAS for Africans were downloaded from GWAS Catalog (https://www.ebi.ac.uk/gwas/home) with catalog number of 37669986 (TC, TG, and HDL-C), 38104120 (DBP and SBP), and GCST90013466 (Height). ROSMAP resources can be requested at https://www.radc.rush.edu. THSBC and CBMAP resources are available on reasonable request from the corresponding author.

## Code availability

The algorithm of MABM model training and association results have been deposited on https://github.com/zdangm/MABM_protein.

## Acknowledgements

This research is supported by the National Natural Sciences Foundation of China 82204118 (D.Z.) and the Healthy Zhejiang One Million People Cohort K-20230085 (D.Z.).

## Author information

## Contributions

D.Z., X.P., and W.Z. designed the study. W.Z. and D.Z. wrote the manuscript. W.Z., L.S., W.F., C.C., and Y.L. collected and processed the data. W.Z. performed the analyses. E.R.G., Y.Z., J.J., L.W. and Y.D. commented, edited and revised the manuscript. D.Z. supervised and acquired funding for the study. D.Z., X.P., and A.P. verified the underlying data. All authors have read and approved the final version of the manuscript.

## Ethics declarations

## Competing interests

The authors declare no competing interests.

## Notes

### Competing Interest Statement

The authors have declared no competing interest.

## References

1. Jiang, L. et al. A Quantitative Proteome Map of the Human Body. Cell 183, 269–283.e19 (2020).

2. Suhre, K., McCarthy, M. I. & Schwenk, J. M. Genetics meets proteomics: perspectives for large population-based studies. Nat Rev Genet 22, 19–37 (2021).

3. Wang, D. et al. A deep proteome and transcriptome abundance atlas of 29 healthy human tissues. Molecular Systems Biology (2019) doi:10.15252/msb.20188503.

4. Lundberg, E. et al. Defining the transcriptome and proteome in three functionally different human cell lines. Molecular Systems Biology (2010) doi:10.1038/msb.2010.106.

5. Bai, Z. et al. Integrating plasma proteomics with genome-wide association data to identify novel drug targets for inflammatory bowel disease. Sci Rep 14, 16251 (2024).

6. Ou, Y.-N. et al. Identification of novel drug targets for Alzheimer’s disease by integrating genetics and proteomes from brain and blood. Mol Psychiatry 26, 6065–6073 (2021).

7. Wingo, T. S. et al. Brain proteome-wide association study implicates novel proteins in depression pathogenesis. Nat Neurosci 24, 810–817 (2021).

8. Gamazon, E. R. et al. A gene-based association method for mapping traits using reference transcriptome data. Nat Genet 47, 1091–1098 (2015).

9. Gusev, A. et al. Integrative approaches for large-scale transcriptome-wide association studies. Nat Genet 48, 245–252 (2016).

10. Brandes, N., Linial, N. & Linial, M. PWAS: proteome-wide association study—linking genes and phenotypes by functional variation in proteins. Genome Biol 21, 173 (2020).

11. Wingo, A. P. et al. Integrating human brain proteomes with genome-wide association data implicates new proteins in Alzheimer’s disease pathogenesis. Nat Genet 53, 143–146 (2021).

12. Zhang, J. et al. Plasma proteome analyses in individuals of European and African ancestry identify cis-pQTLs and models for proteome-wide association studies. Nat Genet 54, 593–602 (2022).

13. Zeng, B. et al. Multi-ancestry eQTL meta-analysis of human brain identifies candidate causal variants for brain-related traits. Nat Genet 54, 161–169 (2022).

14. Xu, F. et al. Genome-wide genotype-serum proteome mapping provides insights into the cross-ancestry differences in cardiometabolic disease susceptibility. Nat Commun 14, 896 (2023).

15. Geoffroy, E., Gregga, I. & Wheeler, H. E. Population-Matched Transcriptome Prediction Increases TWAS Discovery and Replication Rate. iScience 23, (2020).

16. Molstad, A. J. et al. Heterogeneity-aware integrative regression for ancestry-specific association studies. Biometrics 80, ujae109 (2024).

17. Tian, P. et al. Multiethnic polygenic risk prediction in diverse populations through transfer learning. Front. Genet. 13, (2022).

18. Mahajan, A. et al. Genome-wide trans-ancestry meta-analysis provides insight into the genetic architecture of type 2 diabetes susceptibility. Nat Genet 46, 234–244 (2014).

19. Gurdasani, D., Barroso, I., Zeggini, E. & Sandhu, M. S. Genomics of disease risk in globally diverse populations. Nat Rev Genet 20, 520–535 (2019).

20. Zhou, D. et al. A unified framework for joint-tissue transcriptome-wide association and Mendelian randomization analysis. Nat Genet 52, 1239–1246 (2020).

21. Kjolby, M., Nielsen, M. S. & Petersen, C. M. Sortilin, Encoded by the Cardiovascular Risk Gene SORT1, and Its Suggested Functions in Cardiovascular Disease. Curr Atheroscler Rep 17, 18 (2015).

22. Gustafsen, C. et al. The Hypercholesterolemia-Risk Gene SORT1 Facilitates PCSK9 Secretion. Cell Metabolism 19, 310–318 (2014).

23. Musunuru, K. et al. From noncoding variant to phenotype via SORT1 at the 1p13 cholesterol locus. Nature 466, 714–719 (2010).

24. Takeichi, M. The cadherin superfamily in neuronal connections and interactions. Nat Rev Neurosci 8, 11–20 (2007).

25. Stranger, B. E., Stahl, E. A. & Raj, T. Progress and Promise of Genome-Wide Association Studies for Human Complex Trait Genetics. Genetics 187, 367–383 (2011).

26. Li, B. & Ritchie, M. D. From GWAS to Gene: Transcriptome-Wide Association Studies and Other Methods to Functionally Understand GWAS Discoveries. Frontiers in Genetics 12, (2021).

27. George, S. H. L., Medina-Rivera, A., Idaghdour, Y., Lappalainen, T. & Romero, I. G. Increasing diversity of functional genetics studies to advance biological discovery and human health. The American Journal of Human Genetics 110, 1996–2002 (2023).

28. Martin, A. R. et al. Clinical use of current polygenic risk scores may exacerbate health disparities. Nat Genet 51, 584–591 (2019).

29. Lu, Z., et al. Multi-ancestry fine-mapping improves precision to identify causal genes in transcriptome-wide association studies. The American Journal of Human Genetics 109, 1388–1404 (2022).

30. Wainberg, M. et al. Opportunities and challenges for transcriptome-wide association studies. Nat Genet 51, 592–599 (2019).

31. Teslovich, T. M. et al. Biological, clinical and population relevance of 95 loci for blood lipids. Nature 466, 707–713 (2010).

32. Luningham, J. M. et al. Bayesian Genome-wide TWAS Method to Leverage both cis- and trans-eQTL Information through Summary Statistics. The American Journal of Human Genetics 107, 714–726 (2020).

33. Watanabe, K. et al. A global overview of pleiotropy and genetic architecture in complex traits. Nat Genet 51, 1339–1348 (2019).

34. Sun, B. B. et al. Plasma proteomic associations with genetics and health in the UK Biobank. Nature 622, 329–338 (2023).

35. Bycroft, C. et al. The UK Biobank resource with deep phenotyping and genomic data. Nature 562, 203–209 (2018).

36. Das, S. et al. Next-generation genotype imputation service and methods. Nat Genet 48, 1284–1287 (2016).

37. Zhou, D., et al. The China Brain Multi-omics Atlas Project (CBMAP). bioRxiv (2025) doi:10.1101/2025.04.30.651445.

38. De Jager, P. L. et al. A multi-omic atlas of the human frontal cortex for aging and Alzheimer’s disease research. Sci Data 5, 180142 (2018).

39. Tibshirani, R. Regression Shrinkage and Selection via the Lasso. Journal of the Royal Statistical Society. Series B (Methodological) 58, 267–288 (1996).

40. Li, S., Cai, T. T. & Li, H. Transfer Learning for High-Dimensional Linear Regression: Prediction, Estimation and Minimax Optimality. Journal of the Royal Statistical Society Series B: Statistical Methodology 84, 149–173 (2022).

41. Sakaue, S. et al. A cross-population atlas of genetic associations for 220 human phenotypes. Nat Genet 53, 1415–1424 (2021).

42. Nam, K., Kim, J. & Lee, S. Genome-wide study on 72,298 individuals in Korean biobank data for 76 traits. Cell Genomics 2, (2022).

43. Feng, Y.-C. A. et al. Taiwan Biobank: A rich biomedical research database of the Taiwanese population. Cell Genomics 2, (2022).

44. Chen, C.-Y. et al. Analysis across Taiwan Biobank, Biobank Japan, and UK Biobank identifies hundreds of novel loci for 36 quantitative traits. Cell Genom 3, 100436 (2023).

