## Supplemental Data 1 for "Cross-ancestry information transfer framework improves protein abundance prediction and protein-trait association identification"

### Supplementary Figures

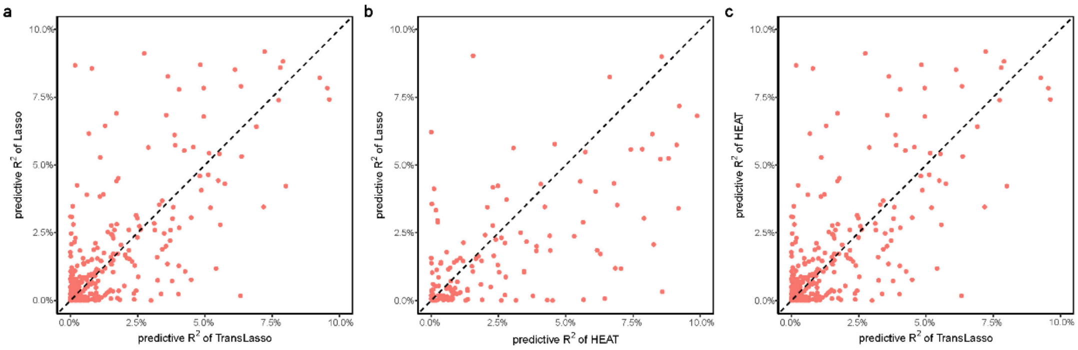


**Supplementary Fig. 1:** Comparison of predictive $R^2$ in THSBC test data between (**a**) TransLasso and Lasso, (**b**) HEAT and Lasso, (**c**) TransLasso and HEAT. Models with a predictive $R^2 \leqslant 1\%$ in the 5-fold cross-validation are not included. Each circle represents a protein.

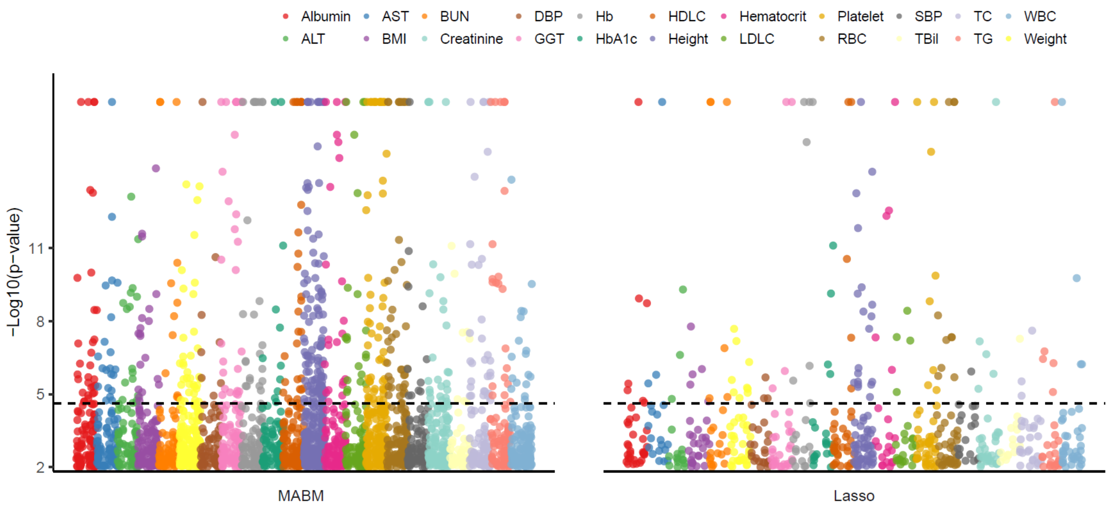


**Supplementary Fig. 2: MABM substantially increased the identification of protein-trait associations using meta-GWAS summary data.** Based on the GWAS summary statistics from meta-GWAS, PWAS was conducted for 22 complex traits using MABM (left) and the baseline Lasso model (right), respectively. The Manhattan plot shows the association signals. The dashed line represents the Bonferroni-significant level (*P*-value < $2.38\times10^{-5}$ and *P*-value < $8.49\times10^{-5}$, respectively) for the two models. The 22 traits of meta-GWAS were labeled with different colors.

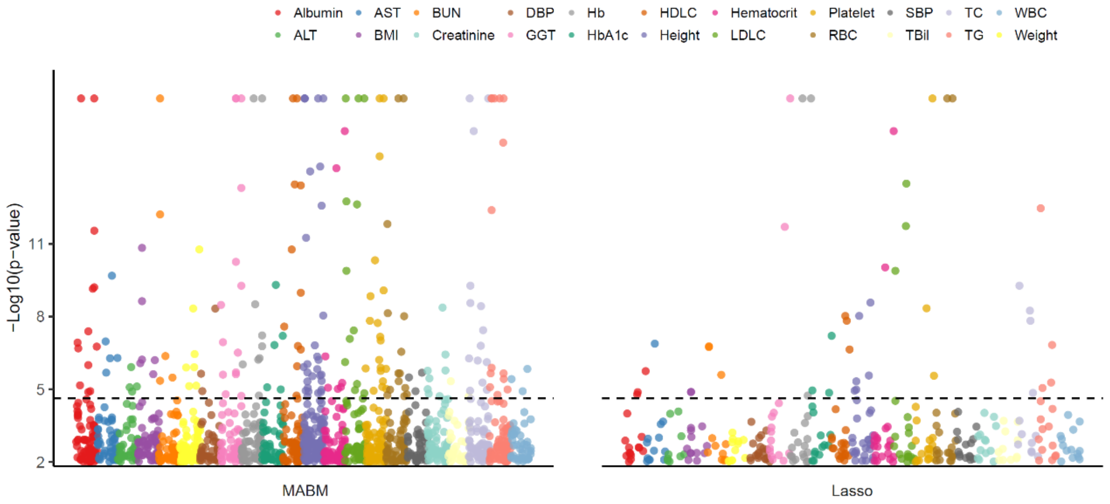


**Supplementary Fig. 3: MABM substantially increased the identification of protein-trait associations in KoGES.** Based on the GWAS summary statistics from KoGES, PWAS was conducted for 22 complex traits using MABM (left) and the baseline Lasso model (right), respectively. The Manhattan plot shows the association signals. The dashed line represents the Bonferroni-significant level ($P$-value < $2.38\times10^{-5}$ and $P$-value < $8.49\times10^{-5}$, respectively) for the two models. The 22 traits of KoGES were labeled with different colors.

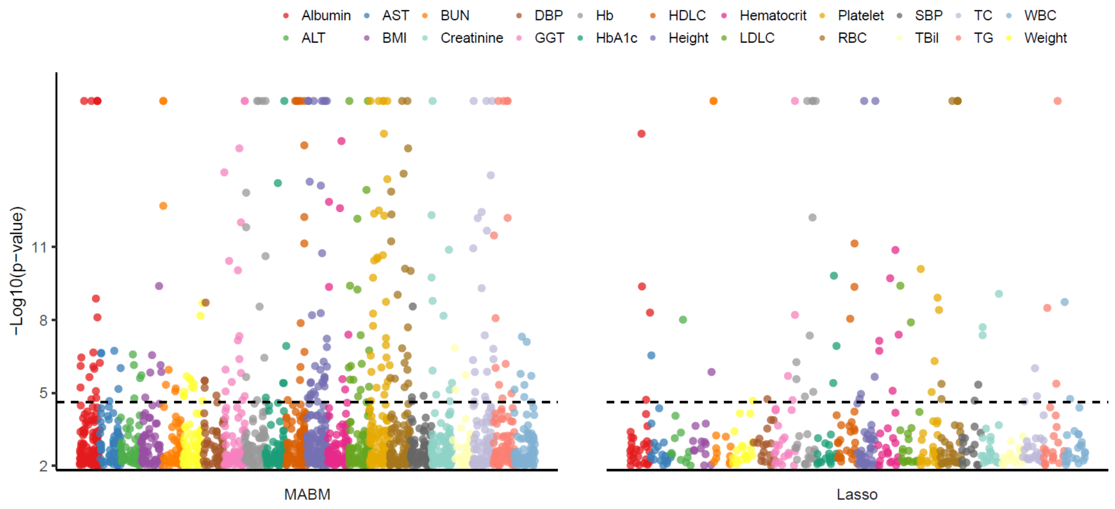


**Supplementary Fig. 4: MABM substantially increased the identification of protein-trait associations in TWB.** Based on the GWAS summary statistics from TWB, PWAS was conducted for 22 complex traits using MABM (left) and the baseline Lasso model (right), respectively. The Manhattan plot shows the association signals. The dashed line represents the Bonferroni-significant level ($P$-value < $2.38\times10^{-5}$ and $P$-value < $8.49\times10^{-5}$, respectively) for the two models. The 22 traits of TWB were labeled with different colors.

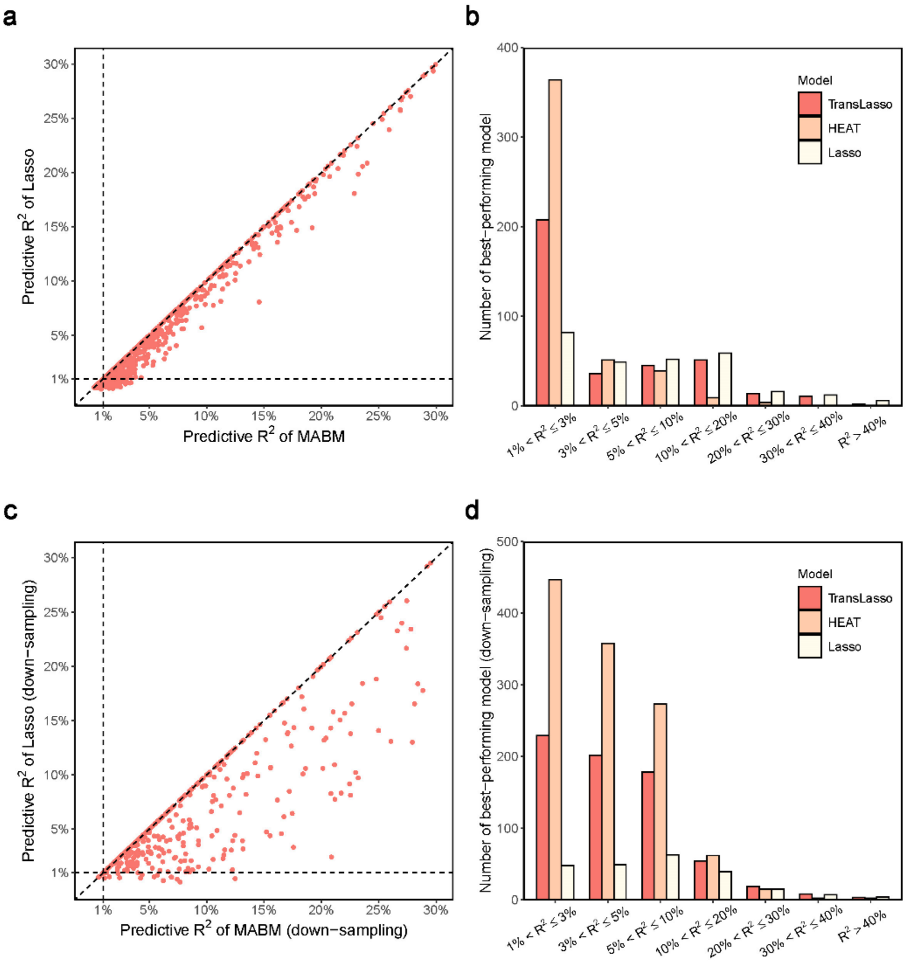


**Supplementary Fig. 5: Comparison of model performance in cross-validation in the AFR populations.** Panel **a** and **c** compare the predictive $R^2$ between MABM and Lasso. Each circle represents a protein. The horizontal and vertical dashed lines represent predictive $R^2$ = 1%. The diagonal dashed line y = x is also presented. Panel **b** and **d** display the composition of best-performing models (MABM) sourced from different model training approaches, partitioned by prediction performance $R^2$. The first row presents the results from cross-validation using all 931 AFR samples, and the second row shows the results from the down-sampling analysis.

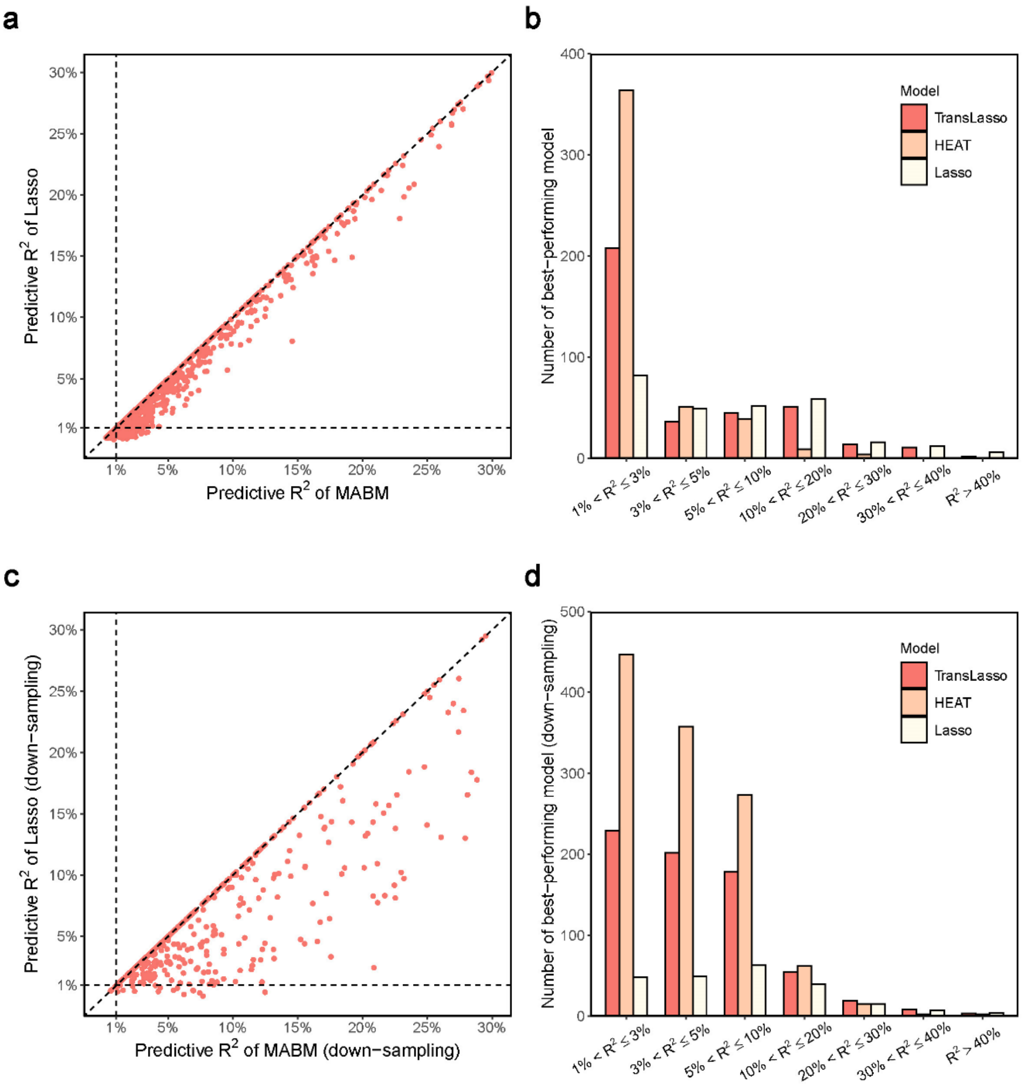


**Supplementary Fig. 6: Comparison of model performance between TransLasso and Lasso in cross-validation across different ancestries.** Each scatter plot represents a comparison within a specific population: (**a**) EAS (*n* = 264), (**b**) MID (*n* = 311), (**c**) CSA (*n* = 923), and (**d**) AFR (*n* = 931). Each circle represents a protein. The horizontal and vertical dashed lines represent predictive $R^2$ = 1%. The diagonal dashed line y = x is also presented.

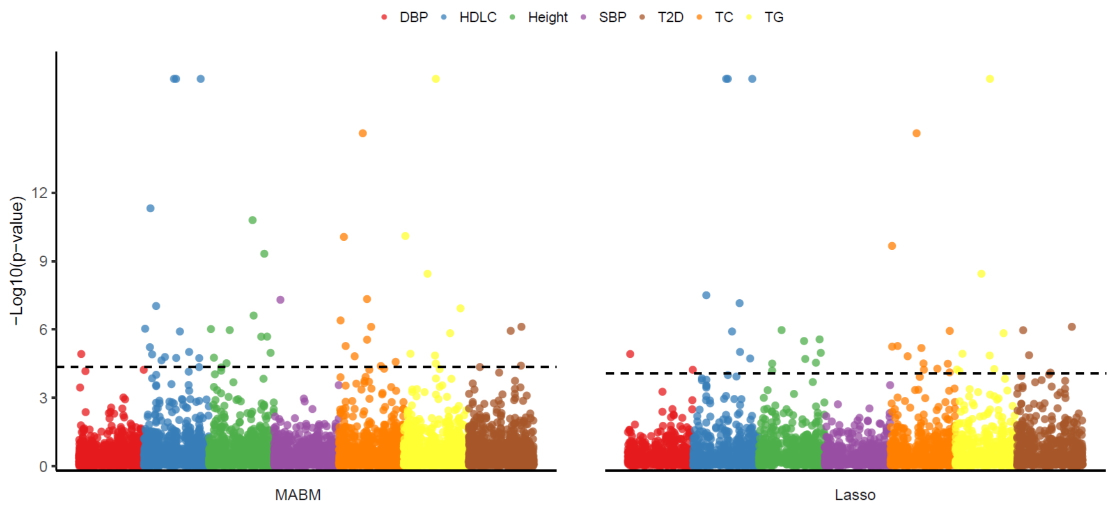


**Supplementary Fig. 7: MABM substantially increased the identification of protein-trait associations in AFR.** Based on the GWAS summary statistics from AFR, PWAS is conducted for seven complex traits using MABM (left) and the baseline Lasso model (right), respectively. The Manhattan plots display the association signals detected by each model. The dashed lines represent the Bonferroni-significant level ($P$-value < $4.52\times10^{-5}$ for MABM and $P$-value < $8.20\times10^{-5}$ for Lasso, respectively). The seven traits are labeled with different colors.

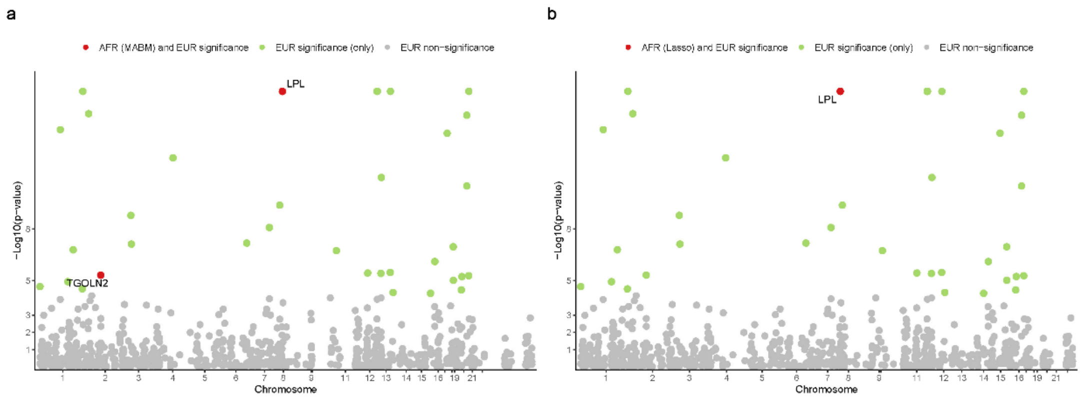


**Supplementary Fig. 8: The integrated PWAS results for HDL-C across EUR and AFR.** Panel **a** presents the Manhattan plot of PWAS results for HDL-C in EUR. We further decorate the results from MABM-based PWAS in AFR. Red, green, and gray circles denote proteins with both AFR and EUR significance, only EUR significance, and EUR non-significance, respectively. The proteins with both AFR and EUR significance are displayed using *P*-values in EUR PWAS and are labeled. Panel **b** presents the same results except for using the Lasso prediction model in AFR PWAS.

### Supplementary Tables

**Supplementary Table 2.** MABM-based PWAS can identify more significant proteins than Lasso-based PWAS across different datasets

| Dataset | MABM-based PWAS | Lasso-based PWAS | Shared identifications | Unique identifications by MABM-based PWAS |
|---|---|---|---|---|
| BBJ | 320 | 101 | 65 | 255 |
| KoGES | 207 | 54 | 31 | 176 |
| TWB | 269 | 84 | 46 | 223 |
| Meta-GWAS | 595 | 147 | 86 | 509 |

### Supplementary Methods

#### Cross-validation setting

In our study, the predictive $R^2$ was evaluated through a nested 5-fold cross-validation[1–3] framework. Below, we provide a more detailed description of our cross-validation procedure:

1. Data splitting: The EAS dataset was randomly partitioned into five equal-sized subsets.
2. Training and validation: In each fold, for Lasso model training, four subsets (80% of the EAS data) were used as the training set, while the remaining subset (20% of the EAS data) served as the test set. For TransLasso model training, four EAS subsets and all sample data from other ancestries were used as the training set, while the remaining EAS subset served as the test set. For HEAT model training, four EAS subsets and all European sample data were used as the training set, while the remaining EAS subset served as the test set.
3. Hyperparameter selection: Within the training set, an inner cross-validation was performed to determine the optimal hyperparameters for each model (Lasso, TransLasso, and HEAT). More specifically, the training dataset was divided into five equal parts, 80% of which was used to train the model and 20% of which was used to test the model performance. Then, the hyperparameters that can achieve the highest prediction accuracy were obtained. In this step, the Lasso model was implemented using the cv.glmnet() function in R, and TransLasso[4] and HEAT[5] were implemented using their internal function settings.
4. Model evaluation in each of the test set: Using the optimized parameters, the model was retrained on the entire training set and then applied to the held-out test set to compute the predictive $R^2$, which was defined as the square of the Pearson correlation coefficient between the predicted and actual protein abundances of the test set.
5. Final performance estimation: This process was repeated five times, ensuring that each subset served as the test set exactly once. The average predictive $R^2$ across all five test sets was taken as the final evaluation metric for each model.
6. Imputable protein definition: Proteins with a predictive $R^2$ greater than 1% were classified as imputable proteins (iProteins).
7. Best model selection: For each protein, we selected the model with the highest $R^2$ as the best-performing model (i.e., MABM). The selected model for each protein had an $R^2 >$ 1% in the nested five-fold cross validation.

### Supplementary Results

##### MABM surpasses Lasso in non-European ancestries with limited sample sizes

We trained TransLasso, HEAT, and Lasso models using genomic and proteomic data from 931 African ancestry individuals in the UKB-PPP dataset. In the five-fold cross-validation, the cross-ancestry models increased the number of iProteins (i.e., the number of models with predicted $R^2 > 1\%$) from 613 (Lasso) to 941 (HEAT) and 765 (TransLasso). However, pairwise comparisons of predictive $R^2$ for shared iProteins showed that the best-performing model identified (i.e., MABM) exhibited a limited improvement over Lasso (Supplementary Fig. 5a,b). Despite this, MABM successfully generated predictive models for 1,110 iProteins, with the median predictive $R^2$ increasing from 2.34% to 2.88% for the less predictable iProteins ($R^2 < 5\%$ in the baseline Lasso model).

Regarding the observation that there is a significant improvement in EAS ancestry ($n = 264$) but limited improvement in African ancestry ($n = 931$), we hypothesize that this may be related to the sample size of non-European ancestry populations. To test our conjecture, we first evaluate it by down-sampling the African ancestry samples. We randomly selected 200 African ancestry individuals and repeated the five-fold cross-validation. The number of iProteins obtained was 437 for Lasso, 1,205 for TransLasso, and 1,701 for HEAT. We obtained MABM models for 2,077 iProteins, and the median predictive $R^2$ increased from 2.80% to 3.96% for the less predictable iProteins ($R^2 < 5\%$ in the baseline Lasso model). The MABM outperformed Lasso just like the condition we observed for EAS ancestries (Supplementary Fig. 5c,d).

Based on the results of down-sampling, we speculate that when the sample size of non-European ancestries is small, sharing information from other populations significantly improves the predictive accuracy of the model. However, when the sample size of non-European ancestries reaches a certain scale, such as 1,000 cases, non-European ancestries themselves can provide a relatively sufficient sample size to model the contribution of genetic variants while also capturing potential ancestry-specific regulation. At this point, the assistance from European ancestry information becomes very limited. This finding underscores the crucial role of cross-ancestry information sharing in small-sample settings, while in large-sample scenarios, single-ancestry models tend to exhibit greater stability and precision.

In addition, we verified our hypothesis using empirical data by evaluating the performance of the trans-ancestry model across multiple non-European ancestry populations. The results showed that, compared to the lasso model, the most significant improvements in prediction performance were observed in the following order: EAS ($n = 264$), MID ($n = 311$), CSA ($n = 923$), and AFR ($n = 931$). We found that the magnitude of improvement was inversely proportional to the sample size of non-European ancestries (Supplementary Fig. 6).

Since the improvement in prediction performance with MABM is relatively limited, we applied both MABM and Lasso models to seven GWAS studies and observed a reasonable improvement. At a Bonferroni-significant threshold of 0.05 ($P$-value < $4.52\times10^{-5}$ for MABM and $P$-value < $8.20\times10^{-5}$ for Lasso), 49 and 42 significant associations were found by MABM and Lasso model, respectively (Supplementary Fig. 7). This indicates that even limited improvements can help us detect more signals, although the magnitude of the increase is not as pronounced as in EAS. It should be underscored that the MABM-based association test achieved an equal or more significant level than the Lasso-based ones in nearly all (95.7%) protein-trait associations identified by both models. This further demonstrates that the increased number of protein-trait associations identified by MABM is not simply due to an increase in the number of iProteins, but rather stems from enhanced prediction accuracy and the resulting boost in statistical power.

We then integrated the PWAS results of HDL-C in Africans with those in Europeans to improve the ability to identify potential causal genes by combining cross-ancestry information. Here, a standard protein abundance imputation model for EUR ancestry was trained using the Lasso model based on the European proteome data from the UKB-PPP. PWAS was performed by applying the EUR model to the HDL-C GWAS summary statistics from UKB European ancestry samples. Its results were presented alongside the PWAS results obtained from MABM using African HDL-C meta-GWAS summary data on the same Manhattan plot (Supplementary Fig. 8a). We identified 35 HDL-C-associated proteins in EUR ($P$-value < $6.01\times10^{-5}$). Among the 35 proteins, the AFR PWAS using MABM probed two proteins *LPL* and *TGOLN2* ($P$-value < $4.52\times10^{-5}$). In contrast, the Lasso model only found one protein (*LPL*, $P$-value < $8.20\times10^{-5}$) overlapping with the proteins identified in EUR (Supplementary Fig. 8b). *LPL* is a recognized risk gene for HDL-C[6–8]. Evidence showed that the microsomal triglyceride transfer protein (*MTP*) signal colocalized with the trans-Golgi network (*TGN*) marker *TGOLN2* (also known as *TGN38*), which has triglyceride transfer activity in Golgi apparatus-rich fractions from mouse liver[9], suggesting that *TGOLN2* may be involved in lipid metabolism. Other studies have shown that *TGN* may be involved in the transport of cholesterol in cells[10–12].

In summary, we showed that MABM surpasses Lasso in non-European ancestries with limited sample sizes. Moreover, MABM dynamically adapts to the features of empirical data (such as sample size) based on its actual performance, offering the flexibility to reduce to the baseline model when necessary. Simultaneously, the enhanced prediction performance achieved through multi-ancestry information sharing can be translated into the association test, reflected not only in the increased number of identified associations but also in the strength of the signals.

#### Supplementary References


1. Stone, M. An Asymptotic Equivalence of Choice of Model by Cross-Validation and Akaike's Criterion. *Journal of the Royal Statistical Society: Series B (Methodological)* **39**, 44–47 (1977).
2. Varma, S. & Simon, R. Bias in error estimation when using cross-validation for model selection. *BMC Bioinformatics* **7**, 91 (2006).
3. Cherlin, S., Howey, R. A. J. & Cordell, H. J. Using penalized regression to predict phenotype from SNP data. *BMC Proc* **12**, 38 (2018).
4. Li, S., Cai, T. T. & Li, H. Transfer Learning for High-Dimensional Linear Regression: Prediction, Estimation and Minimax Optimality. *Journal of the Royal Statistical Society Series B: Statistical Methodology* **84**, 149–173 (2022).
5. Molstad, A. J. *et al.* Heterogeneity-aware integrative regression for ancestry-specific association studies. *Biometrics* **80**, ujae109 (2024).
6. Nishiwaki, M. *et al.* Effects of alcohol on lipoprotein lipase, hepatic lipase, cholesteryl ester transfer protein, and lecithin:cholesterol acyltransferase in high-density lipoprotein cholesterol elevation. *Atherosclerosis* **111**, 99–109 (1994).
7. Jin, W., Marchadier, D. & Rader, D. J. Lipases and HDL metabolism. *Trends in Endocrinology & Metabolism* **13**, 174–178 (2002).
8. Jin, W., Millar, J. S., Broedl, U., Glick, J. M. & Rader, D. J. Inhibition of endothelial lipase causes increased HDL cholesterol levels in vivo. *J Clin Invest* **111**, 357–362 (2003).
9. Swift, L. L. *et al.* Subcellular localization of microsomal triglyceride transfer protein. *Journal of Lipid Research* **44**, 1841–1849 (2003).

10. Maxfield, F. R. & Wüstner, D. Intracellular cholesterol transport. *J Clin Invest* **110**, 891–898 (2002).

11. Urano, Y. *et al.* Transport of LDL-derived cholesterol from the NPC1 compartment to the ER involves the trans-Golgi network and the SNARE protein complex. *Proceedings of the National Academy of Sciences* **105**, 16513–16518 (2008).

12. Reverter, M. *et al.* Cholesterol Regulates Syntaxin 6 Trafficking at trans-Golgi Network Endosomal Boundaries. *Cell Reports* **7**, 883–897 (2014).
